# p53motifDB: integration of genomic information and tumor suppressor p53 binding motifs

**DOI:** 10.1101/2024.09.24.614594

**Authors:** Gabriele Baniulyte, Sawyer M. Hicks, Morgan A. Sammons

## Abstract

The tumor suppressor gene *TP53* encodes the DNA binding transcription factor p53 and is one of the most commonly mutated genes in human cancer. Tumor suppressor activity requires binding of p53 to its DNA response elements and subsequent transcriptional activation of a diverse set of target genes. Despite decades of close study, the logic underlying p53 interactions with its numerous potential genomic binding sites and target genes is not yet fully understood. Here, we present a database of DNA and chromatin-based information focused on putative p53 binding sites in the human genome to allow users to generate and test new hypotheses related to p53 activity in the genome. Users can query genomic locations based on experimentally observed p53 binding, regulatory element activity, genetic variation, evolutionary conservation, chromatin modification state, and chromatin structure. We present multiple use cases demonstrating the utility of this database for generating novel biological hypotheses, such as chromatin-based determinants of p53 binding and potential cell type-specific p53 activity. All database information is also available as a precompiled sqlite database for use in local analysis or as a Shiny web application.

## INTRODUCTION

Sequence-specific transcription factors are key regulators of cellular and developmental processes (Lambert et al., 2018). Understanding the mechanisms by which transcription factors accurately recognize and bind to cognate DNA motifs and discriminate against other DNA sequences has been a central question in molecular and developmental biology for decades (Jolma et al., 2013). The number of potential transcription factor (TF) motifs in the genome far outweighs the number of observed binding events (Inukai et al., 2017), suggesting that sequence alone does not fully dictate binding. Transcription factors must also integrate multiple other types of DNA and chromatin-embedded information during binding site selection *in vivo.* Nucleosome positioning, histone and DNA modifications, and chromatin conformation join DNA sequence and shape as factors regulating transcription factor binding to DNA (Neph et al., 2012; Thurman et al., 2012; Wang et al., 2012). For example, the majority of transcription factors cannot bind to their cognate motif when the sequence is engaged with a nucleosome providing one molecular mechanism reducing the ratio of observed to potential binding events (Zaret and Carroll, 2011; Thurman et al., 2012).

Areas of open chromatin flanked by nucleosomes with particular histone modification patterns can be used as indirect evidence for regulatory region activity (Bannister and Kouzarides, 2011; Klemm et al., 2019; Millán-Zambrano et al., 2022). Massive efforts in mapping genomic locations of open chromatin and histone modification localization revealed clear patterns of cell type specificity in transcriptional regulation (Dorschner et al., 2004; Bernstein et al., 2005; Kundaje et al., 2015). These data have been successfully integrated with transcription factor binding information and other data sources, such as genetic variation and evolutionary conservation, to provide insight into the biological function and mechanisms of transcriptional regulatory regions (Wang et al., 2012; Vierstra et al., 2020).

The tumor suppressor gene *TP53* encodes a DNA binding transcription factor called p53. The well-characterized tumor suppressor activity of p53 requires two distinct but related functions: DNA binding and the activation of transcription (Kastenhuber and Lowe, 2017). DNA binding is central to the activity of p53, as the majority of cancer-associated *TP53* mutations are found in the DNA binding domain (DBD) and disrupt interactions between p53 and its cognate DNA response element (Hollstein et al., 1991). The nucleotide preferences within a p53 response element/motif (p53RE) have been rigorously validated using multiple low and high-throughput methodologies, including EMSA, SELEX, and ChIP-seq (el-Deiry et al., 1992; Wei et al., 2006; Jolma et al., 2013). DNA sequence variation across p53REs only partially explains observations of differential p53 binding and transcriptional activity *in vivo* (Wang et al., 2009; Vyas et al., 2017), suggesting that other genomic features may serve a key role in these processes. Recent work suggests that p53:DNA interactions differ between cell types (Nguyen et al., 2018a), and are influenced by features such as local and long distance chromatin structure (Serra et al., 2024), histone modifications (Isbel et al., 2023), and DNA methylation (Kribelbauer et al., 2017). Differential p53 binding is itself linked to differential p53 transcriptional targets across cell types, suggesting that p53 activity can be modulated by differences in cell or condition-specific chromatin structure.

In this paper, we present the p53motifDB, which integrates predictions of p53 response element motifs within the human genome with multiple genetic, epigenetic, and functional datasets. The goals of the p53motifDB are 1) to act as a comprehensive resource for quickly obtaining key information about both putative and validated p53 binding events in the human genome and 2) to serve as a tool to generate novel hypotheses about p53-dependent transcriptional regulation. The entire database is available as standalone tables for integration into machine learning paradigms or as a precompiled sqlite database. Users can also access the p53motifDB via a local or web-based shiny App. We provide multiple examples of the utility of integrating these datasets by confirming previous observations in the field and by generating and testing novel hypotheses.

## MATERIALS AND METHODS

### Data sources and processing

Table 1 contains the file name, data type, source/download location, and relevant publication (if applicable) for each dataset used to construct the p53motifDB. Putative p53 motifs in the GRCh38/hg38 genome assembly were identified using two separate methodologies. We first used p53 motifs defined by JASPAR (matrix ID MA0106.3) from an HT-SELEX-derived position weight matrix (PWM) (Jolma et al., 2013). We then used a PWM derived from experimental ChIP-seq data and the GRCh38/hg38 assembly as the inputs for running the scanMotifsGenomewide.pl script from HOMER (Heinz et al., 2010). The output of the two p53 motif datasets were merged using bedTools to create a non-redundant master list of 412,586 individual motifs (Quinlan and Hall, 2010). The liftOver tool and corresponding chain files were used to identify corresponding loci in the hg19 and T2T/hs1 human genome assemblies and to identify syntenic locations in the mm10 and MM39 mouse genome assembly (Hinrichs et al., 2006). Unless otherwise stated, p53 motif locations were integrated with datasets available in interval formats using bedTools. Data extraction from bigWig file types was performed using deepTools (Ramírez et al., 2016). The dbSNP156 dataset in vcf file format was queried using bcftools/samtools (Sherry et al., 1999, 2001). Data were parsed from program-specific output files and joined into database-compatible tables using *dplyr* and *tidyr* packages implemented in *R* (version 3.6.0) (Wickham et al., 2018; Wickham and Henry, 2019).

### Data access and Shiny web app construction

Processed datasets were added into a SQLite3 relational database built using the DBI (v.1.2.3) and RSQLite (v.2.3.7) packages within *R* (version 3.6.0). The database primary key is the hg38 genome coordinate for each motif in the “chr_start_stop” format, called the “unique_id” in each table. SQLite3 database files are available to download from Zenodo (10.5281/zenodo.13351805). We also built a Shiny app under R (version 3.6.0) that allows users to access the database via an online portal (https://p53motifdb.its.albany.edu/). The Shiny app and code can be downloaded for off-line use from Zenodo or can be deployed locally via a pre-built Docker image (Merkel, 2014; Chang et al., 2024). All raw data tables used to construct the SQLite3 database and the Shiny app can also be downloaded directly from Zenodo under CC-BY data restrictions.

## RESULTS

### Characteristics and selection of p53 response elements (p53RE) in the human genome

The content of the p53motifDB is focused on providing key genetic and regulatory information on non-redundant genomic locations that contain putative p53 binding locations based on experimentally validated p53 binding preferences. We thus began our investigation by identifying potential p53 response element motifs (p53RE) in the human genome. The canonical p53 response element motif contains two half-sites separated by a 6 nucleotide spacer (el-Deiry et al., 1992). This concept of identifying potential p53RE in the genome is not new, and multiple algorithms and approaches have been previously used to identify genomic p53RE (Wei et al., 2006; Smeenk et al., 2008; Veprintsev and Fersht, 2008; Cui et al., 2011). All approaches used either *in vitro* or ChIP-style data to derive motif likelihood scores based on p53 affinity for particular DNA sequences. Recent meta-analyses of dozens of ChIP-seq datasets suggest that most *in vivo* p53 binding occurs at “canonical” motifs (Nguyen et al., 2018b; Riege et al., 2020). Therefore, we did not consider p53 ½ or ¾ sites or potential motifs with spacers which would greatly increase the number of potential p53RE with only a marginal gain in *bona fide* binding events (el-Deiry et al., 1992). We allowed for overlapping p53RE, with the 3’ half-site of one p53RE serving as the 5’ half-site of another. Our approach depended on updated p53RE PWM from high-throughput *in vitro* (SELEX) and *in vivo* (ChIP-seq) and two separate software approaches. We first extracted pre-compiled p53 motif locations from hg38 in the JASPAR 2022 database (Castro-Mondragon et al., 2022) which uses SELEX data to define the PWM. These motifs were merged with putative p53RE identified using scanMotifGenomeWide.pl from HOMER (v4.10.4) (Heinz et al., 2010) and its built-in p53 position weight matrix (p53.motif) derived from ChIP-seq data.

We then standardized the length of all p53RE from HOMER and JASPAR to 20 nucleotides to account for the different lengths of the underlying PWM used to call motif locations, and converted all motif locations to the plus-strand. This resulted in a total of 412,586 non-redundant p53 response element (p53RE) motifs on the canonical somatic (1-22), sex (X/Y) and mitochondrial (MT) chromosomes (Fig. 1A). Motifs present on unassigned scaffolds were removed from further analysis. Interestingly, only 113,858 motifs were identified by both JASPAR and HOMER (Fig. 1A). The JASPAR database contained substantially more unique p53RE (273,678) than those identified by HOMER (25,050) (Fig. 1A). p53RE identified in both datasets are scored higher (i.e more closely aligned to the consensus) than those identified via HOMER (Fig. 1B) or JASPAR (Fig. 1C) alone. p53RE identified uniquely by HOMER had nearly identical preferences for the key C/G residues within the half-sites to shared sites (Fig. 1D vs. Fig. 1E), whereas unique JASPAR elements had reduced prevalence for these crucial nucleotides (Fig. 1D vs. Fig. 1F). HOMER-specific motifs were depleted for purines at the 5’ end and pyrimidines at the 3’ end relative to p53RE shared across methods (Fig. 1E). The nucleotide frequency differences between HOMER and JASPAR (Fig. 1E vs. Fig. 1F) may reflect either differences in the statistical methods used to call motifs or differences in experimental approaches used to derive the underlying PWM (*in vitro* SELEX vs. chromatin immunoprecipitation). We also marked p53RE that contain potential cytosine methylation sites (6.9%, 28,485/412,586). Biochemical and structural evidence indicates that DNA methylation within a p53RE can strongly influence p53 binding and transcriptional regulatory activity (Kribelbauer et al., 2017). Methylation at CG dinucleotides within a p53RE will be cell- and context-dependent and should be considered as part of any comprehensive analysis of p53 biochemical activity on DNA.

**Figure 1:**
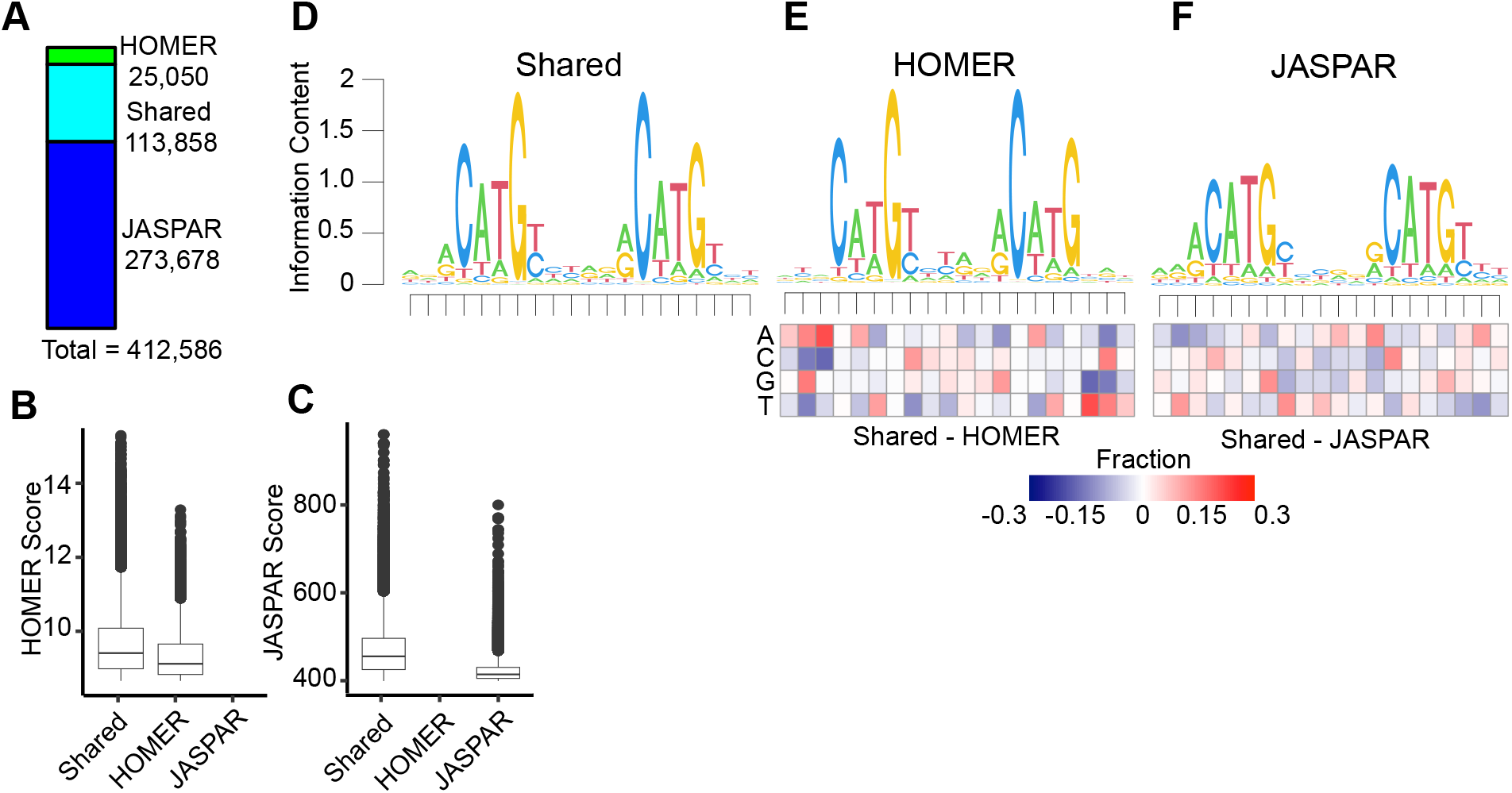
Selection and characteristics of p53 response elements (p53RE) in the hg38 human genome assembly. **A)** The number of p53RE identified using nucleotide binding preferences from HOMER (ChIP-seq-derived) compared to JASPAR (*in vitro* SELEX-derived). The distribution of method-specific scoring for p53RE either found with both methods or with only a single method for **B)** HOMER or **C)** JASPAR. **D)** A seqLogo representation of nucleotide preferences for p53RE identified from both HOMER and JASPAR binding preferences. seqLogo representation of nucleotide preferences for **E)** HOMER-specific or **F)** JASPAR-specific p53RE. Each heatmap represents the nucleotide frequency between the shared p53RE and either the HOMER or JASPAR-specific p53RE.

### Integration with other reference genome assemblies and repetitive elements

In order to extend the utility of these datasets, we examined whether the standardized 412,586 p53RE motifs and their locations were also present in two additional human reference genome assemblies and in two commonly-used mouse reference genomes using the UCSC liftOver tool (Hinrichs et al., 2006; Kuhn et al., 2013). Providing information from additional mouse and human reference genomes will allow users to more quickly integrate p53-centric information with the wealth of pre-parsed data from prior assemblies (hg19 and mm10). Further, the inclusion of genomic locations from the recently completed full telomere-to-telomere human hs1 genome will provide some level of future-proofing as new datasets are provided with updated genomic coordinates (Nurk et al., 2022). As expected, nearly all hg38 p53RE locations are present in the hg19 genome assembly and are preserved in the most recent hs1/T2T complete human genome assembly (>99%, Table 2).

We also report p53RE intersections with two additional key pieces of genome assembly data. First, we incorporated genome blacklist information into p53motifDB which covers 6,163 locations in the hg38 genome. These locations are generally excluded from most genomic analyses due to pervasive issues in mapping next-generation sequencing reads from experiments like ChIP-seq and RNA-seq to a reference genome assembly (Amemiya et al., 2019). We also provide information on the location of p53RE relative to repetitive DNA elements. Repetitive DNA elements, like those derived from transposable elements and viral DNA, significantly contribute to difficulty in genome assembly and next-generation sequencing-based analyses of genome function (Treangen and Salzberg, 2012). Repetitive elements also influence gene regulatory networks (Feschotte, 2008; Sundaram et al., 2014), serve as platforms for transcription factor binding (Bourque et al., 2008) and can have *cis*-regulatory element activity (Chuong et al., 2017). p53 is known to bind to and regulate activity of repetitive elements (Wang et al., 2007, p. 53; Harris et al., 2009; Tiwari et al., 2018). We provide information regarding the presence of p53RE within repetitive DNA elements using the RepeatMasker compendium (Jurka, 2000; Smit et al., 2013). Over 60% of identified p53RE (254,075/412,586) are found in repetitive elements. In addition to whether a p53RE is located within a p53RE, our dataset includes information about the repeat DNA itself, including the repeat class and family.

Mouse models have been extensively used to identify foundational tenets of p53 biology. Tumor suppressor activity is conserved across vertebrates, but p53 biochemical activity and specific genes regulated by p53 differs between mouse and human (Stewart-Ornstein et al., 2017; Fischer, 2019). Functional transcription factor binding events controlling gene expression often vary between even closely related organisms (Tanay et al., 2005). Prior work identified a limited number of p53RE with conserved sequence and binding across a range of vertebrates with a focus on whether these p53RE might be functionally linked to gene expression (Horvath et al., 2007; Jegga et al., 2008). We extend this work here by examining synteny between p53RE locations in the human genome and two commonly-used mouse genome assemblies. As expected, the majority of p53RE from hg38 are unique to human assemblies and are not syntenic within the mouse genome. We found approximately 21% of p53RE locations are syntenic in the MM10 (88,624) or MM39 (89,074) mouse reference genomes. Within the database, we provide the genomic coordinates from the hg38 assembly and, if applicable, the corresponding coordinates in the hg19, hs1/T2T, MM10, or MM39 genomes. We’ve also provided an average phyloP and phastCon vertebrate conservation scores for the 20bp p53RE (Pollard et al., 2010; Siepel et al., 2005) allowing the end user to rapidly identify human p53 motifs with different evolutionary conservation constraints. Ultimately, the conservation and synteny data presented here are not meant to replace a more comprehensive and focused evolutionary analysis of p53RE, but can be used in combination with other datasets as a means to quickly generate and test hypotheses related to the p53 gene regulatory network.

### Integration and analysis of p53 genomic binding datasets

The p53motifDB also contains information on whether individual p53 motifs are locations for experimentally observed p53 binding as determined by four recent meta-analyses of dozens of p53 chromatin immunoprecipitation-coupled sequencing (ChIP-seq) datasets (Verfaillie et al., 2016; Nguyen et al., 2018b; Riege et al., 2020; Hammal et al., 2022). Overall, 37,628 p53RE have evidence of p53 binding from ChIP-seq data in at least one cell line or condition (Fig. 2). There are 3,833 p53RE with evidence of experimental p53 binding across all 4 meta-analyses (Fig. 2). Each meta-analysis used a different combination of p53 ChIP-seq datasets, different genome mapping methods, and statistical approaches for calling positive binding events, resulting in a range of binding events across meta-analyses. Each meta-analysis only considered p53 binding events that met some statistical or enrichment cut-off based on the peak calling tool of choice. With the exception of the ReMAP dataset, each meta-analysis also only considered a p53 binding event as legitimate if observed across multiple experiments or cell lines. The Verfaillie dataset characterized both “Strong” and “Weak” p53 binding sites from 16 individual p53 ChIP-seq datasets from a total of 7 different cell types based on inferred affinity from read pileup data. We combined both Strong and Weak categories into one group to simplify our analysis, yielding 4,922 p53RE that overlap p53 binding events. Of note, this analysis considered the smallest number of datasets. Nguyen et al used two different metrics for determining p53 binding based on the number of datasets in which a given putative binding site reached a predetermined cutoff value (Nguyen et al., 2018a). We use the nomenclature “ubiquitous” and “validated” from the original publication to maintain consistency across analyses. These analyses yielded a set of highly-stringent, “ubiquitous” p53 binding events (1,288 sites, identified in >= 20 independent datasets) and a less-stringent set of “validated” (12,048 sites, >= 2 independent datasets) p53 binding sites. We used only the less-stringent set of p53 binding events in this analysis. Riege et al used an experimental cut-off of 5 independent datasets, resulting in 7,804 p53RE with observed p53 occupancy. Finally, the ReMap2022 dataset considered 126 p53 ChIP-seq datasets, and included all binding events that occurred in any individual experiment. In total, 3,833 p53RE are occupied by p53 across the Nguyen (validated), Verfaillie, Riege, and Remap datasets. An additional 3,654 p53RE have p53 binding in at least three of these datasets (Fig. 2). The Remap dataset contains 24,322 p53 ChIP-seq binding events not found in any other meta-analysis, likely due to the combination of a larger number of datasets considered and a reduced threshold for calling a binding event relative to other datasets. Differences between datasets also represent tissue and cell-specific binding events, including the Nguyen dataset which identified cell lineage-specificity to p53 binding (Nguyen et al., 2018a). Cell type-specificity for p53 binding has been further confirmed in additional studies (Nguyen et al., 2018b; Karsli Uzunbas et al., 2019; Hafner et al., 2020). The majority of human p53 genomic binding data comes from transformed cell lines, thus we expect that the number of p53RE with observed p53 binding events will likely increase as additional cell lineages, primary cells and tissues, and as new stimulus paradigms are considered. The full dataset includes genomic coordinates for each observed ChIP-seq peak and experimental observation frequency data in supplemental database tables for ReMap and Riege meta-analyses, and included information about the cell lines used for analysis from the ReMap2022 dataset.

**Figure 2:**
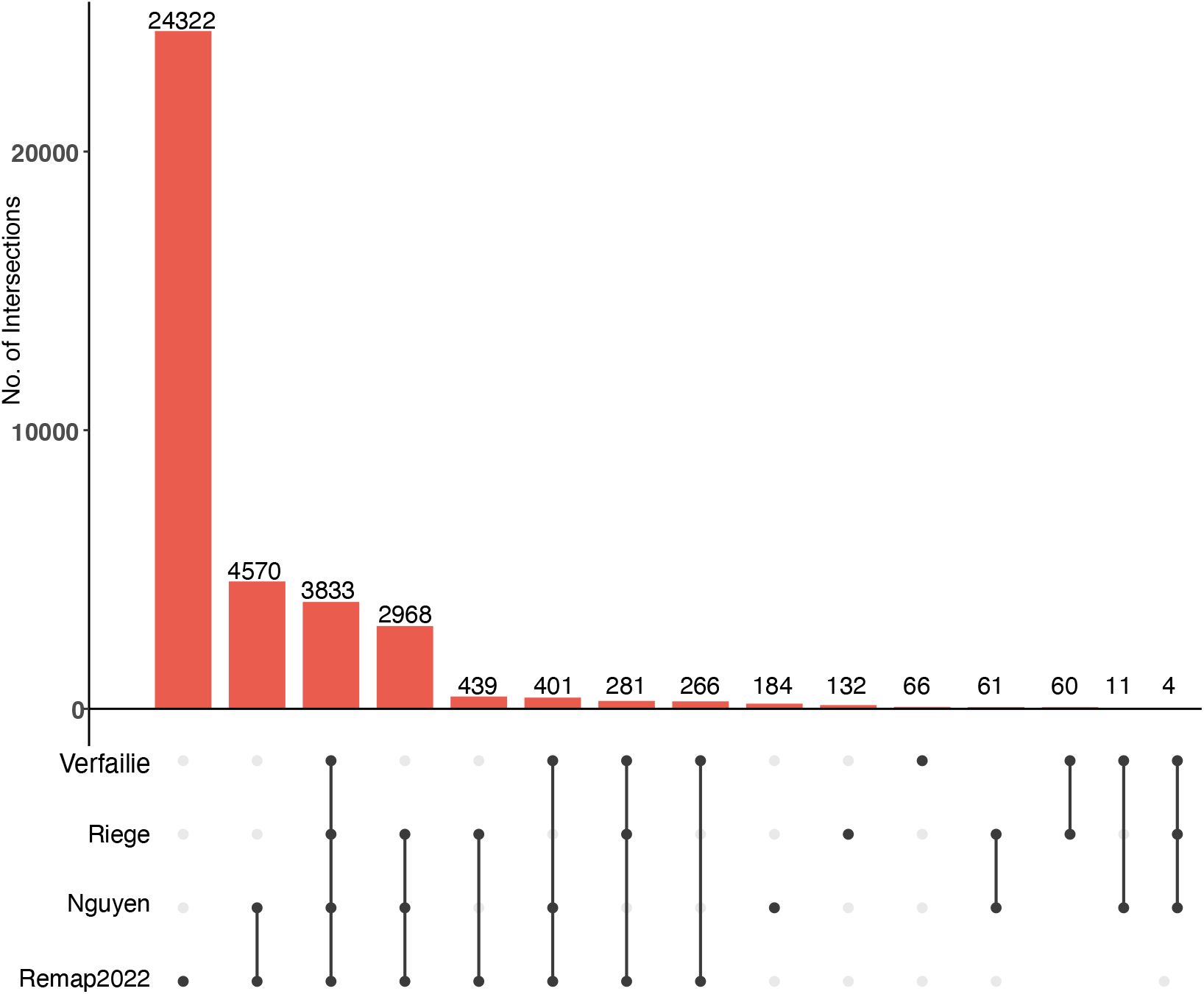
Overlap between identified p53RE and experimentally-validated p53 binding events across four comprehensive meta-analyses. Upset-style plot of overlap between p53RE and experimentally-derived p53 binding events (via ChIP-seq) for four separate meta-analyses.

### Integration of gene features and gene expression changes with p53RE locations

p53 is a transcription factor regulating a diverse set of target genes under multiple physiological conditions, including in response to DNA damage. We included a number of datasets to help researchers explore relationships between the location of p53REs and gene expression. Most p53RE are found within ENSEMBL gene models (intragenic, 61%, 253,727/412,586) compared to outside genes (intergenic, 39%, 158,859/412,586). When applicable, the HGNC gene symbol is also provided (i.e *TP53*). The dataset can be filtered by HGNC symbol, ENSEMBL gene ID (ENSG*) or ENSEMBL transcript ID (ENST*) when those values are available. The frequency of all intragenic p53RE is nearly identical to that of bound p53RE (61% vs. 63%) when considering the 3,833 p53RE that are occupied by p53 across all four ChIP-seq meta-analyses (Fig. 2). These data suggest there is no innate preference for actual p53 binding within genes other than the natural distribution of p53RE across the genome. We also report distance to the nearest transcriptional start site both up and downstream of the p53RE, and include ENSEMBL gene ID, transcript ID, and HGNC gene symbols associated with those TSS. We also integrated gene expression fold-change data and gene expression “scores” from a prior meta-analysis of p53’s effect on transcription (Fischer et al., 2022). Thus, researchers can rapidly search the database for genes and transcripts whose activity is known to be influenced by p53 and identify the nearest set of p53RE and p53 binding events. We note that linking TF binding to gene expression changes is a complex process that requires more than correlation between proximity of two elements, but that this information can be used to generate hypotheses for downstream functional validation.

### Integration of local chromatin and regulatory element information with p53RE

Local chromatin structure strongly influences transcription factor binding and activity (Neph et al., 2012; Thurman et al., 2012). Multiple publications have examined how these features can influence p53 interactions with the genome and subsequent transcriptional activation (Sammons et al., 2015; Su et al., 2015; Verfaillie et al., 2016). We thus incorporated multiple summary datasets describing chromatin modification states and chromatin accessibility into the database to allow further exploration of these potential linkages. DNAse I hypersensitive sites (DHS) are genomic regions susceptible to enzymatic cleavage, thus reflecting “open”, nucleosome-free locations (Vierstra and Stamatoyannopoulos, 2016). We incorporated ENCODE DHS cluster data which integrates millions of DHS locations across 125 cell types and conditions (Sheffield et al., 2013; Moore et al., 2020). These data include the cell type where the DHS was observed and an accessibility score based on the maximum observed DNAse signal. Overall, 86,851 p53REs (21%) are found within DHS. The ENCODE Project also produced a series of analyses focused on predicting regions of the genome that function as candidate Cis-Regulatory Elements (cCRE) based on local chromatin structure and gene distance (Moore et al., 2020). The analysis produced 5 cCRE classes: promoter (high DNase and H3K4me3 signal, <200bp from TSS), promoter-like (high DNase and H3K4me3 signal, >200bp from TSS), proximal enhancer (high DNase and H3K27ac, low H3K4me3, <2kb from TSS), distal enhancer (high DNase and H3K27ac, low H3K4me3, >2kb from TSS), and CTCF-binding (high CTCF and DNase, low H3K4me3 and H3K27ac). A total of 48,177 p53RE (11.67% of total) overlap an ENCODE cCRE. Within that 48,177, 2.5% are within the promoter group, 2.0% are promoter-like, 12.8% are at proximal enhancer elements, 79.6% at distal enhancers, and 3.0% overlap CTCF-binding elements.

Chromatin accessibility (DNase hypersensitivity/ATAC-seq) can be combined with multiple chromatin modification states to predict transcription regulatory activity. The ENCODE and Epigenome Roadmap projects produced thousands of histone modification datasets across a vast array of cell and tissue types (The ENCODE Project Consortium, 2012; Kundaje et al., 2015). While these data provided incredible insight into connections between the epigenome, transcription factors, and gene expression, the breadth and scope of these datasets is often unwieldy for non-computational biologists. ChromHMM uses hidden Markov models to incorporate multiple types of chromatin modification and accessibility data to summarize the likely functional chromatin state of a given genomic location across different cell types (Ernst et al., 2011; Ernst and Kellis, 2015, 2017). An updated fullstack chromHMM segmentation was designed to create a “single universal” chromatin-based annotation for different segments of the genome (Vu and Ernst, 2022). Thus, we integrated the “fullstack” chromHMM genome segmentation dataset in order to simplify analysis of local chromatin context surrounding p53RE. The dataset includes 100 detailed chromatin-state annotations which can be collapsed into 16 broader chromatin state “groups” as previously defined (Vu and Ernst, 2022). Nearly all p53RE are located within a chromHMM genome annotation segment (99%, 409,061/412,586), providing researchers with a simple description of the most-likely local chromatin environment for a given p53RE.

### Incorporation of three-dimensional chromatin interactions with p53RE locations

Assigning individual transcription factor binding events to specific gene expression changes can be difficult without additional genetic evidence, such as the deletion of a specific transcription factor binding site or regulatory element. Connecting TF binding and gene activity is difficult even when it occurs at a gene promoter, and is even more difficult for distal TF binding events (Chen et al., 2020). Chromatin looping has emerged as a key driver of transcription and partially explains how distal regulatory elements can control gene expression over long distances (Krivega and Dean, 2012; Kulaeva et al., 2012; Dekker et al., 2013). Here, we integrate four datasets interrogating three-dimensional chromatin interactions with p53RE locations. The GeneHancer and Activity by Contact (ABC) datasets incorporate chromatin conformation assays with gene expression and regulatory element-associated activity data, like chromatin modifications, to call enhancer:promoter interactions across a range of cell types and conditions (Fishilevich et al., 2017; Fulco et al., 2019). Promoter capture approaches use *in situ* proximity-based ligation approaches to identify distal regions interacting with targeted promoters. This database incorporates recently published promoter-capture Hi-C focused on p53-mediated gene regulation from HCT116 cells (Serra et al., 2024) and newly generated promoter-capture Micro-C (GEO GSE275042) from MCF10A mammary epithelial cell lines treated with either DMSO or the p53 activating drug etoposide. Two important caveats should be considered when assessing Hi-C-based approaches and their application to p53 biology. First, the experimental and statistical bias in Hi-C analysis is biased towards identification of distal interactions. Since many p53RE are within or near promoters, we expect reduced sensitivity in detecting TSS-proximal interactions. Second, the resolution of the technique (normally 1kb+) means it can be difficult to determine specific p53RE or p53 binding events actually participating in a looping event. To streamline the analysis and standardize the data for use in this database, biological and technical replicates, treatment conditions, and time points for the promoter-capture experiments were merged to create a single “snapshot” of potential 3D interactions involving p53RE. Across all 4 datasets, a total of 206,575/412,586 p53RE (50.06%) are found within at least one chromatin looping interaction. Localization within a chromatin loop anchor does not necessarily imply transcriptional regulatory potential, just as the absence from a loop does not mean a given p53RE is not functional. These data allow users of the p53motifDB to quickly identify and generate hypotheses regarding experimentally-observed chromatin loops containing potential or validated p53 binding events without having to parse or re-analyze large chromatin conformation capture datasets.

### Human genetic variation at p53RE

Reference genomes do not represent the vast diversity of genomic space in the human population. Genetic variation in *cis*-regulatory elements and transcription factor binding sites can directly influence biochemical activities on DNA (Spivakov et al., 2012). Genetic variation in transcription factor binding sites can also lead to a range of additional biological effects, ranging from changes in gene expression up to observable phenotypic traits with the potential to alter health and lifespan. The DNA sequence-based determinants of p53 binding and activity are well-studied (el-Deiry et al., 1992; Wang et al., 2009), but are still under active investigation (Fischer and Sammons, 2024). Variation in p53RE motifs can have strong functional effects *in vivo*. Users may be interested in whether p53 binding and gene regulation might be affected by natural human genetic variation in p53REs. We therefore incorporated genetic variation data from the dbSNP156 build which includes millions of single nucleotide polymorphism and small insertion and deletion variants (Sherry et al., 2001). Over 99% of p53REs have at least one reported genetic variant from dbSNP156 (408,872/412,586), with a median of 9 variants per p53RE. p53RE almost universally contain single nucleotide variants (SNV), but deletions (33%, 136,826/412,586) or insertions (7.8%,32,261/412,586) of at least 1 nucleotide are also relatively common. We’ve also included variation reported in the clinVar database, which contains variants with known or predicted clinical significance (Landrum et al., 2018, 2020). For example, rs4590952 (G>A) is a common SNP in a p53RE that reduces p53 binding, alters transcription of *KITLG,* and is associated with increased cancer risk (Zeron-Medina et al., 2013). The master p53motifDB table contains information only about whether a given p53RE contains a clinVar or other SNP, but that information can be used to query a secondary table containing information about the specific reference and alternate allele for each SNP in the database.

The data integrated into this current database specifically focus on variation within the canonical 20bp p53 motif. Variation outside of this core motif may influence p53 binding and activity, such as variation within other TF binding motifs required for regulatory element activity (Korkmaz et al., 2016; Catizone et al., 2020). Further, our analysis does not consider genetic variation that could result in *de novo* p53 motifs and gene regulation as has been previously observed (Menendez et al., 2007, 2019). The rapid increase in genome resequencing projects focused on human genetic diversity, personal genomes, and cancer genomics should allow these and other types of advanced analyses in the future.

### Accessing and querying the p53motifDB

We have provided multiple methods for users to interface with the data found in the p53motifDB. Our goal was not to produce a comprehensive analysis platform, rather, we aimed to provide end users with multiple, flexible methods of interacting with this dataset. The tables were built and analyzed via *R* using *tidyverse-*style data methods, but users can easily import the underlying data into their preferred data analytics tools and pipelines. All processed datasets are available in tabular-format via Zenodo. A pre-compiled *sqlite* relational database is also fully-available for download via Zenodo which can be analyzed offline using standard query methods. Power users, or those who wish to perform more advanced relational queries are encouraged to download either the sql database or the tabular-format datasets for use in custom data analysis pipelines.

We also integrated the datasets into a Shiny app, which can be queried online (https://p53motifDB.its.albany.edu) or downloaded and used locally from our Zenodo repository (10.5281/zenodo.13351805). The interface for the Shiny app allows users to initially filter the p53motifDB based on categorical information in the main table. Drop down boxes with predefined choices are available for data types with a limited number of choices, such as when searching by chromosome or searching for p53RE with experimentally-observed p53 binding. Input boxes can be used for other types of data, such as when querying by gene names. By default, p53REs are filtered by whether there is experimental evidence of p53 binding in any of the 4 p53 ChIP-seq meta-analyses representing hundreds of assays. Data can be further filtered and results can be exported to a local tab-delimited file for offline analysis. Users can export either the primary table information or can query additional data sources using the filtered p53RE locations. As an example, a user might filter p53RE that are found in an ENCODE DHS cluster and where there is experimental p53 binding evidence, but then want to know more information about the cell types where those DHS are found. Users can quickly export this advanced information from accessory tables, like DHS clusters or genetic variation, based on filtered p53RE locations by selecting one of a series of buttons in the Shiny app. All underlying database information and code for building and deploying the Shiny app are available on the Zenodo repository (10.5281/zenodo.13351805).

### Use cases for the p53motifDB

### Enrichment of p53RE and p53 binding in repetitive elements

Repetitive genomic elements encompass satellite and microsatellite DNA as well as those derived from viral and mobile genetic elements. The contribution of repetitive and mobile genetic elements to the distribution of p53 motifs in the genome is well documented (Tiwari et al., 2018). We identified 254,075 p53RE motifs (61.6%) within repetitive DNA elements using the RepeatMasker UCSC Genome Browser (Jurka, 2000; Smit et al., 2013). Almost half of the p53RE within repetitive elements are contained in LINE elements (43.08%) (Fig. 3A). SINE (32.97%), LTR (13.29%), and DNA (5.31%) and simple (4,73%) repeat elements are also substantial contributors of p53RE motifs (Fig. 3A). All other repetitive element classes add up to approximately 10% of the total. Consistent with the primate specificity of SINE and LINE expansion (Konkel et al., 2010), p53RE found within the MM39 mouse reference genome are considerably less likely than expected to be found in repetitive elements compared to regions lacking synteny (Fig. 3B, *p < 2.2e-16, Pearson’s Chi-squared with Yates’ continuity correction*).

**Figure 3:**
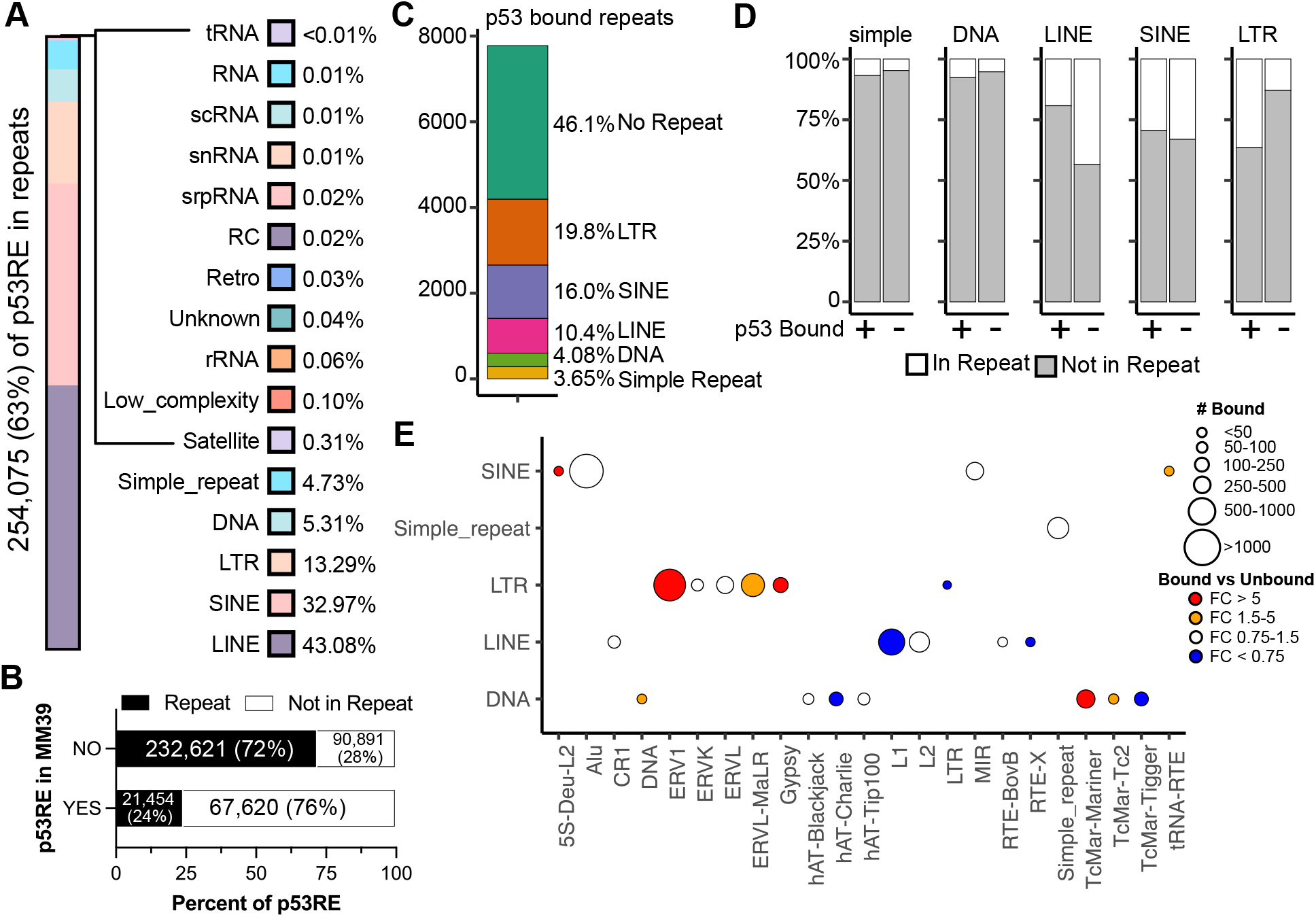
Characterization of p53RE and their localization within repetitive genomic elements. **A)** The percentage of p53RE found within each class of repeat element. A total of 254,075/412,586 p53RE (63%) are found within repeat elements. **B)** The distribution of p53RE with synteny to MM39 and found in repeat elements. **C)** The percentage of p53RE with experimentally-validated p53 binding (Riege et al., 2020) and their distribution within repeat elements. **D)** The distribution of p53-bound versus p53-unbound p53RE and their localization within LINE, SINE, LTR, DNA, and simple repeats, which represent the five most common repeat types with p53RE. **E)** Number of p53-bound p53RE within each class and type of repeat element for the five most common repeat types and their enrichment versus p53-unbound p53RE.

We then asked whether experimentally-determined p53 binding sites are enriched in different repeat element classes, as has been previously demonstrated (Wang et al., 2007; Tiwari et al., 2018). We focused on the p53 binding events from the Riege meta-analysis, as it included a large number of datasets and the threshold for calling a p53 binding event was stringent (i.e p53 binding in at least 5 separate datasets) (Riege et al., 2020). A slight majority of p53 binding events occur within repetitive elements (53.9%), but this is less than expected based on the frequency of repeat-associated p53RE motifs in the genome (61.6%, *p < 2.2e-16, Pearson’s Chi-squared with Yates’ continuity correction*). This suggests actual p53 binding is slightly biased against p53RE within repetitive elements. Compared to p53RE distribution in the genome, p53 binding is significantly enriched within LTR elements (Fig. 3D). ERV1 and ERV1-MaLR elements are preferentially bound relative to the distribution of p53RE genomewide (Fig. 3E). Consistent with prior observations on a more limited set of p53 binding sites, LTR-associated p53 binding frequently occurs within MLT1H, LTR10C/E and MER61C/E elements (Su et al., 2015). In contrast, p53 binding to LINEs is less frequent than expected (Fig. 3D), with L1 LINEs particularly depleted for p53 occupancy and primarily contributing to this observation (Fig. 3E).

### Local chromatin states at p53RE

The local chromatin environment at a given p53 binding site can provide context clues as to whether a particular binding site may function as an enhancer or promoter (Sammons et al., 2015; Su et al., 2015; Serra et al., 2024). Chromatin structure and histone modification patterns at p53 binding sites can differ across cell types which may inform differential activity (Thurman et al., 2012; Nguyen et al., 2018a). We assessed the average chromatin status of experimentally-determined p53 binding sites from the Riege meta-analysis to determine whether p53 is enriched in any specific chromatin locations. First, we assessed the genome-wide enrichment of p53RE in each of the 16 chromHMM summary groups. The distribution of p53RE almost perfectly mirrors the percentage of the genome covered by each chromHMM segment (Fig. 4A).

**Figure 4:**
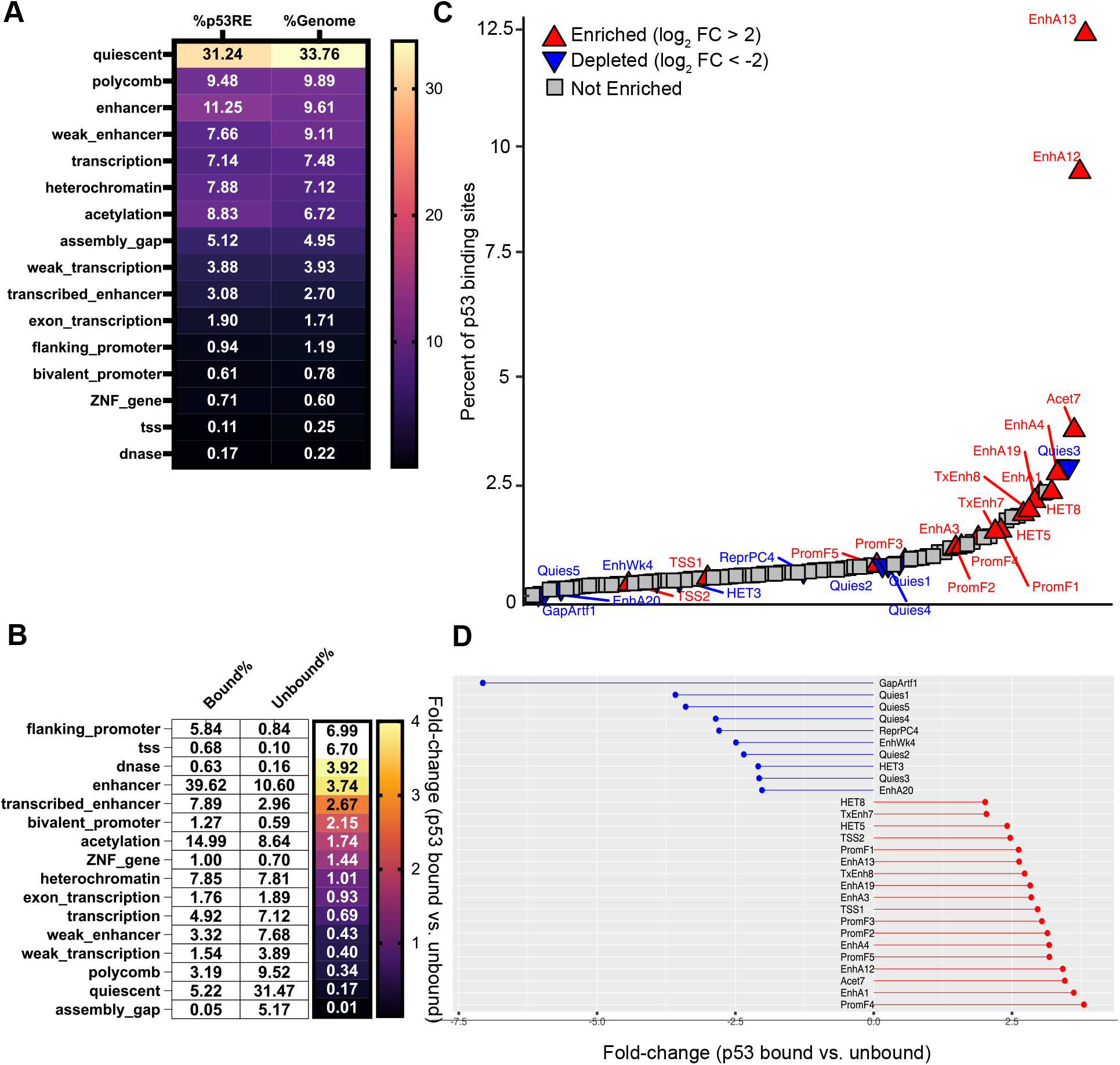
Analysis of p53RE and their localization within chromHMM genome segments. **A)** The percentage of p53RE found within each of the 16 chromHMM fullstack summary groups compared to the percentage of the human genome covered by that chromHMM feature (Vu and Ernst, 2022). **B)** The percentage (white boxes) and fold-change enrichment (color scale) of bound p53RE versus unbound p53RE found within each of the 16 chromHMM fullstack summary groups. p53 binding data are from the Riege et al meta-analysis (Riege et al., 2020). **C)** The rank order of p53-bound p53RE (x-axis) versus the percent of p53-bound p53RE that are found within each of the 100 detailed chromHMM fullstack genomic segments. Arrows represent an enrichment/fold-change of greater than 2 (red upward arrows) or less than -2 (blue downward arrows) for p53-bound versus p53-unbound p53RE. **D)** The actual enrichment/fold-change for p53-bound versus unbound p53RE for each chromHMM fullstack segment labeled in **C.**

We then examined the relationship between chromHMM summary states and p53RE that were bound by p53 versus those that remain unbound. Our analysis suggests that p53 binding is enriched in chromatin contexts reflecting gene regulatory elements, like promoters, enhancers, and other open chromatin regions (Fig. 4B). Nearly 40% of all p53 binding events are found in regions defined as active enhancers, an enrichment of 3.75-fold versus the distribution of unbound p53RE. Promoter and TSS-associated binding was enriched nearly 7-fold. p53 binding is also depleted in regions with H3K27me3/Polycomb-associated heterochromatin, but not in heterochromatin regions enriched with H3K9me3 (Fig. 4B). p53 binding is also depleted strongly from quiescent regions of the genome, which lack chromatin modifications indicative of regulated activity. Combined with the presence of p53 in H3K9me3-marked heterochromatin and the high enrichment in accessible, gene regulatory regions, the lack of p53 binding in quiescent chromatin may be a function of p53’s documented pioneer transcription factor activity (Espinosa and Emerson, 2001; Nili et al., 2010; Laptenko et al., 2011; Sammons et al., 2015; Younger and Rinn, 2017; Yu and Buck, 2019).

We wanted to further demonstrate the utility of integrating p53RE data with local chromatin context by assessing enrichment of p53 binding within sub-categories of chromHMM segments. The fullstack chromHMM dataset contains 100 different chromatin states categorized primarily by enrichment of specific chromatin modifications, cell type, and other genomic features, like gene distance. We graphed these data by the percent of total p53 binding sites in that chromatin segment in rank-order and then color-coded each based on enrichment or depletion status (p53-bound/unbound ratio) (Fig. 4C). Almost 20% of p53 binding events are located in EnhA12 and EnhA13 chromHMM states and p53 binding is enriched over 2-fold relative to unbound p53RE (Fig. 4C-D). EnhA12/EnhA13 represent epithelial-specific enhancer regions, consistent with expanded p53 binding and activity in epithelial cell types (McDade et al., 2014; Karsli Uzunbas et al., 2019; Riege et al., 2020). The PromF4 subclass, representing a chromatin state found downstream of transcriptional start sites, is the most enriched (Fig. 4D). This may reflect the well-documented preference of p53 binding within the first intron of target genes (Zauberman et al., 1995; Takimoto and El-Deiry, 2000; Thornborrow et al., 2002; Liu et al., 2004; Riley et al., 2008). We also observe counterintuitive enrichment of p53 within the HET5 and HET8 subclasses of H3K9me3-heterochromatin (Fig. 4C-D). Although p53 binding was not depleted at H3K9me3-enriched regions like we observed at Polycomb-regulated H3K27me3-enriched regions (Fig. 4B), we did not expect specific enrichment of any subclass of heterochromatin. Interestingly, HET5 and HET8 represent chromatin states found at LTR repetitive elements, which we previously demonstrated support higher-than-expected p53 binding (Fig. 3D). p53 binding is also depleted from the HET3 subclass (Fig. 4C-D) which represents LINE-associated chromatin, consistent with our prior observation of LINE-mediated depletion of p53 binding (Fig. 3D). Taken together, analysis of p53RE and p53 binding enrichment across chromHMM summary and subclass datasets provides further support for prior observations in literature, but also can be used to generate novel hypotheses about p53 activity relative to cell type and chromatin state.

## CONCLUSIONS AND PERSPECTIVE

In summary, we constructed a database resource containing local genetic, regulatory, chromatin, and variation information at putative motifs for the p53 transcription factor in the human genome. These data are accessible in a web-facing application where the end user can query the database without prior knowledge of structured query language. The entire dataset is also available for download in multiple off-line formats allowing more advanced users to analyze the data using tools of their choice. This resource provides users a simple, yet powerful, method for retrieving information on validated p53 binding locations or new putative sites identified in their own laboratories. Finally, we envision this database as a tool that can be used to generate new hypotheses about how chromatin structure, genetic variation, and evolutionary constraint might affect p53 activity across cell types and across organisms with implications in cancer, stem cell, and developmental biology.

## CODE AVAILABILITY

Code and data required to build the sqlite database or the Shiny app can be found in our Zenodo repository (10.5281/zenodo.13351805)

## AUTHOR CONTRIBUTIONS

Project conceptualization: GB, MAS

Data processing: GB, SMH, MAS

Database construction: MAS

Data Generation: GB

Shiny app design: SMH, MAS

Manuscript writing and editing: GB, SMH, MAS

## ACKNOWLEDGEMENTS

We would like to thank all of the labs and research programs that generated the raw and analyzed datasets and metadata used in the construction of this database. We would also like to thank the University at Albany Information Technology Services (ITS) and the Advanced Research Computing Cluster (ARCC) for computing resources, storage, and hosting of the Shiny app.

